# FilmArray Respiratory Panel Integrated in a field Point Of Care dispositive for the diagnosis of respiratory tract infections in rural areas in Senegal

**DOI:** 10.1101/502575

**Authors:** Hubert Bassene, Sophie Edouard, Georges Diatta, Jean Christophe Lagier, Oleg Mediannikov, Florence Fenollar, Aldiouma Diallo, El Hadj Ba, Didier Raoult, Philippe Parola, Michel Drancourt, Cheikh Sokhna

## Abstract

The development of molecular syndrome-based kits for the diagnosis of respiratory infections offers rapid and sensitive detection of common respiratory pathogens and will have a significant impact on the care of patients. In this study, we present the results obtained after the introduction of the FilmArray respiratory panel in a field Point of care (POC) for the diagnosis of virus and bacteria responsible of respiratory tract infections in Senegal rural area. From February to August 2017, we collected nasal swabs from febrile patients that presented symptoms of respiratory tract infections in three health posts located in Niakhar. Specimens were tested on site by multiplex Polymerase Chain Reaction (PCR), using the FilmArray respiratory panel^®^ that targets 20 pathogens, including 17 virus and 3 bacteria (bioMérieux). Nasal swabs were collected from 113 patients. The median age was 4 years (ranging from 4 months to 60 years) and 51 (45%) were males. The prevalence of respiratory pathogens was 37.5% (12/32) during the dry season and 54.3% (44/81) in the rainy season (p=0.16). The prevalence of respiratory pathogen carriage was higher in children under 5 years of age (38/55, 69.1%). The most prevalent micro-organisms detected were influenza B virus (16/113, 14%), human rhinovirus/enterovirus (10/113, 9%), parainfluenzae virus (9/113, 8%), respiratory syncytial virus (8/113, 7%), adenovirus (5/113, 4%), human metapneumovirus (3/113, 3%), *Chlamydia pneumoniae* (2/113, 2%) and *Coronavirus* (2/113, 2%). The study has demonstrated that the integration of the FilmArray respiratory panel into a field POC could significantly improve the management of respiratory tract infections in rural areas.

**Author summary:** Respiratory tract infections are one of the leading causes of death worldwide. Populations in underdeveloped countries are the most affected, especially children under 5 years of age. Among the pathogens responsible for these infections, bacteria and viruses are the most identified. In poor countries, laboratory diagnosis can only be done in urban areas. They are generally based on bacteriological, immunological or molecular biology techniques. The concept of POC is not developed in underdeveloped countries. Its existence would have made it possible to carry out RDTs to confirm the clinician’s diagnosis and allow rapid management of the patient. In the current situation, the time required to achieve results is often very long for patients. In this study, we sought to demonstrate the significant contribution of the FilmArray respiratory panel in the management of respiratory tract infections in rural areas. We also wanted to reduce the time it takes to deliver results in order to improve patient care. This dispositif was integrated in a field POC which was implemented in Niakhar since 2015, in order to improve the management of emerging diseases. The FilmArray respiratory panel gave us the opportunity to investigate the causes of respiratory tract infections in this area. For each patient, we systematically target 20 pathogens, including 17 viruses and 3 bacteria, with a single multiplex PCR. One of the main results of this study is that children under 5 years of age are the most affected by respiratory tract infections. Then we noted a lack of consultation among adults that could be explained by the banalization of respiratory problems or a preference for traditional care. The fact that children under 5 years of age are the most affected could also serve as a basis for implementing vaccination programmes directly targeting the most vulnerable age groups. It should be noted that for 51% of patients, the result of the diagnosis was negative. It would appear that some pathogens responsible for respiratory tract infections are not targeted by the multiplex PCR. Thus, it would be necessary to screen these pathogens in order to integrate them into a panel that would cover the most pathogens in circulation in this area.

## Introduction

Infectious diseases represent a major public health problem in the world. In the poorest countries, they continue to deeply impact economies and paralyze health systems. Progress remains uneven, and millions of people do not benefit from prevention and treatment. In 2013, respiratory infections accounted for just over 4 million deaths worldwide (1). Children under age 5 are most affected by these diseases (2), and lower respiratory tract infections are the most lethal transmittable diseases in these children, with just over 3 million deaths worldwide in 2015 (1). Many bacteria and viruses can cause respiratory tract infections. However, the majority of cases registered has viral cause (3). The most commonly identified viruses are rhinoviruses, coronaviruses, influenza viruses, respiratory syncytial virus (RSV), parainfluenza virus (PIV), human metapneumovirus (hMPV), and adenovirus (4). In most underdeveloped countries, the diagnosis of respiratory infections is based only on clinical signs and symptoms and the absence of laboratory analyzes may have negative consequences on the management of respiratory tract infections including the risk of misdiagnosis. The burden for the population is important because the management of respiratory diseases unrelated to the influenza virus is very expensive. Its direct cost is estimated at $ 17.3 Billion (5).

Several diagnostic methods could be used for the identification of suspected respiratory pathogens. For a long time, it mainly relied on culture but during the last decades, the development of qPCR has revolutionized diagnostic strategies and largely contributed to expand the knowledge on the epidemiology of respiratory diseases, especially for viruses (6). PCR is a method that has a good sensitivity and specificity and allows the concomitant detection of several microorganisms by multiplexing. In addition, the delivery time of results is shorter than for other techniques (7). The best approach could be to target several pathogens in a single run, in order to reduce detection costs, to increase the speed of the diagnosis and finally to target a wide range of pathogens. The devices that are currently available meet many of these requirements. Their differences are in the number of pathogens included in the panel, the species targeted, integrated nucleic acid extraction and the duration of the PCR.

The multiplex PCR BioFire FilmArray Respiratory Panel (bioMérieux) targets 20 pathogens including 17 viruses and 3 bacteria. In this study, we integrated the device into a molecular laboratory Point Of Care (POC) (8) set up in rural areas in Senegal to identify pathogens responsible for febrile respiratory diseases.

## Methods

### Study site

Niakhar is located in the department of Fatick, capital of the region of the same name, and covers 230 km^2^. The study area currently covers a total of 30 villages, where the French National Research Institute for Sustainable Developement (IRD) has been performing an epidemiological and environmental demographic monitoring over the last 50 years. The population of the study site was estimated to be of 44,828 inhabitants in 2017, including 22,095 men and 22,733 women. The area is rural, but the three largest villages are urbanized, with health facilities, weekly markets, several shops and buses that provide daily connection with Dakar, the capital of the country. Health care is provided by three health posts (Toucar, Diohine and Ngayokhem) belonging to two different health districts (Fatick and Niakhar). In 2017, the rainy season started in June and ended in September (IRD data, 2017).

### Samples collection

Nasal swabs were collected from February to August 2017 in febrile patients with a temperature superior or equal to 37.5°C and presenting with signs of respiratory infections attending health posts in Toucar, Diohine and Ngayokhem. We included patients presenting with fever and cough, expectoration, dyspnea, sore throat and/or cold/runny nose. These people systematically benefited from a rapid diagnosis test (RDT) that allowed the detection of malaria, in accordance with the procedures for the management of pathological episodes currently in effect in Senegal. Then the project proposed, in addition to TDR and molecular diagnosis of bacteremia, a nasal swabbing in order to investigate the viruses and bacteria causing respiratory infections.

Samples were taken with VIROCULT swabs (Medical Wire, Corsham, Wiltshire, UK) targeting the inner nasal walls by rotation, in order to collect cells. Samples were conserved as indicated by the manufacturer. Finally, the tube was closed, placed in a cooler at 4°C and sent to the laboratory for analysis. The following age categories (<1, 1-5, 6-21, 22-65 and >65 years) were adopted to study the prevalence rates.

The project was approved by the National Ethics Committee N ° 00081 MSAS / DGS / DS / CNERS. Written informed consent was obtained from all individuals, including patients and parents or legal guardians of all children.

### BioFire FilmArray^®^ description

The diagnosis of respiratory pathogens was performed by a multiplex qualitative PCR, using the BioFire FilmArray^®^ respiratory panel (bioMérieux, France). This method targets 20 respiratory pathogens including 17 viruses and 3 bacteria. The BioFire FilmArray^®^ is placed in a mobile, simple, secure and connected laboratory workstation named POCRAMé^®^. Molecular analyzes performed with this device are automated and simplified, limiting operator intervention.

All reagents required for sample analysis are preloaded in lyophilized form in the reagent storage compartments of the FilmArray^®^ cassette. After loading the hydration solution and the sample to be analyzed, the FilmArray^®^ cassette is loaded into the instrument and the sample number is recorded using a bar code reader.

### Molecular analysis

The FilmArray^®^ instrument integrates all stages of PCR analysis including sample lysis, nucleic acid extraction, amplification, detection and analysis. Approximately 300μL of each sample was used for the diagnosis. The preparation of sample required 2 minutes and a total analysis time of approximately one hour. In this study, the FilmArray^®^ cassettes and instrument were prepared and used by following the manufacturer’s instructions (9, 10).

### Data analysis

Statistical analyses were performed with the Epi Info software 7.0.8.0 version (Centers for Disease Control and Prevention, Atlanta, GA, USA). Data were compared with the Pearson chi2 or Fisher exact tests when applicable and with the statistical significant threshold set at P value ≤ 0.05.

## Results

From February to August 2017, a total of 573 febrile patients were examined at the three health posts in the study area. Nasal swabs were retrieved from 113 patients (32 specimens in the dry season and 81 in winter) that presented, in addition to fever, symptoms of respiratory tract infections. The median age was 4 years (ranging from 4 months to 60 years) and 51 (45%) were males. The peak of swab collection was obtained between July and August with 56% (63/113) of the samples (Figure 1). At least one respiratory pathogen was identified in 42% (48/113) of patients. Globally, the prevalence of respiratory micro-organisms was higher during the rainy season (54.3%, 44/81) but the difference was not significant (χ^2^ = 1.97, *p*=0.16).

**Figure 1:**
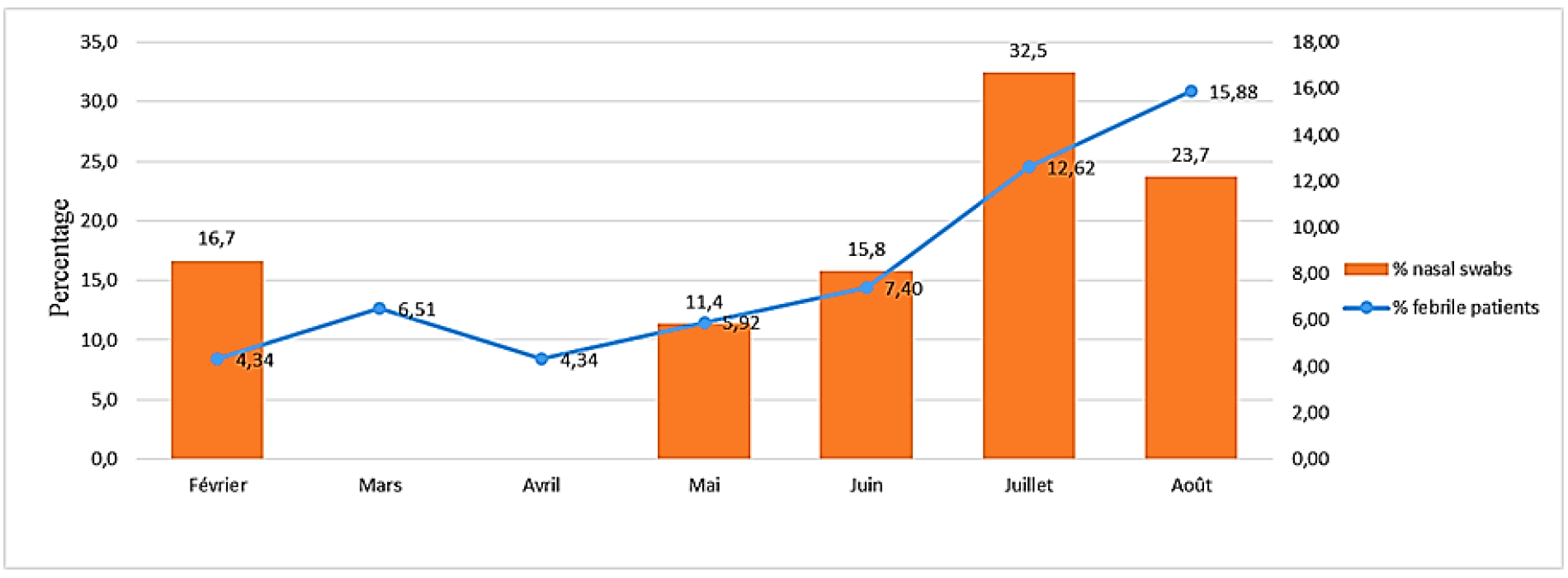
Sample collection during the study period.

The most prevalent micro-organisms detected were influenza B virus (16/113, 14%), human rhinovirus/enterovirus (10/113, 9%), parainfluenzae virus (9/113, 8%), respiratory syncytial virus (8/113, 7%), adenovirus (5/113, 4%), human metapneumovirus (3/113, 3%), *Chlamydia pneumoniae* (2/113, 2%) and *Coronavirus* (2/113, 2%) (Figure 02).

**Figure 2:**
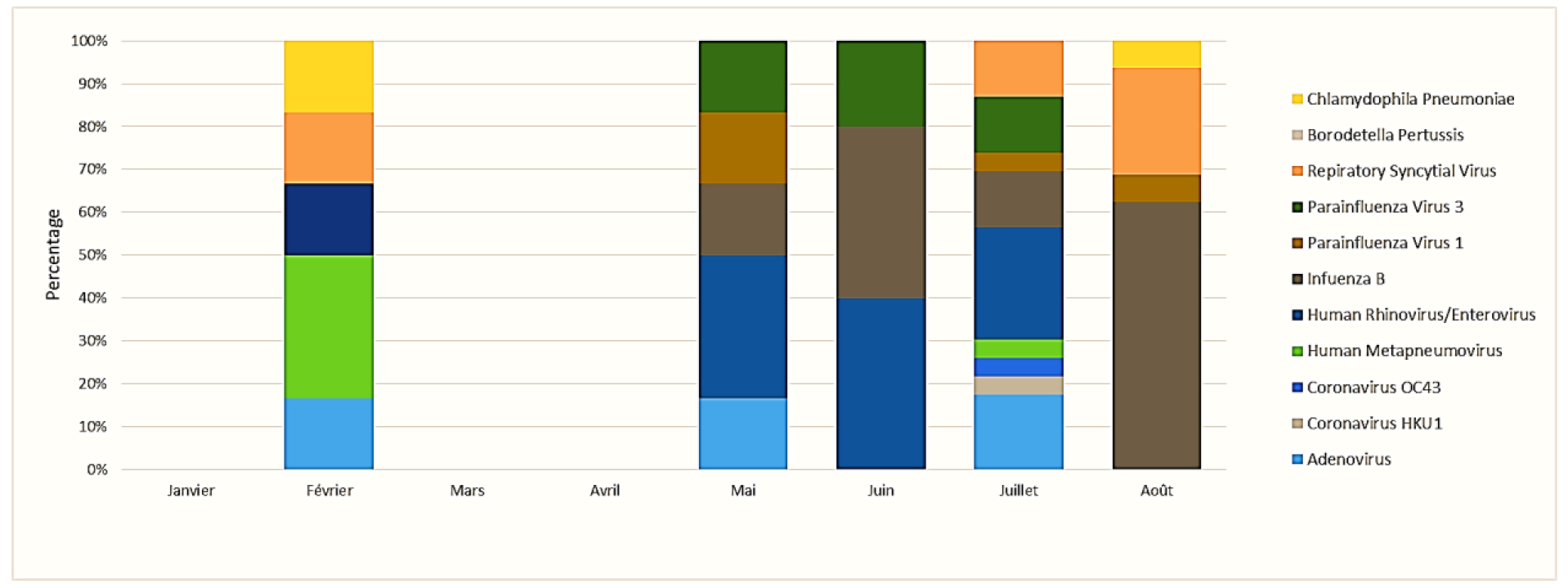
Monthly proportional representation of identified pathogens.

Infections in children under age 5 (38/55, 69.1%) represented the majority of infections. The highest detection rate for respiratory pathogens (11/12, 92%) was noted in infants (< 1 year category), followed by the 1-5 years category (27/49, 55%) (Table 1). The difference between these two categories was significant (χ^2^ = 4.04, *p*<0.05). No pathogen was identified in the 22-65 years category. Infections in the 1-5 category was more diversified, 11 pathogens were identified, more than in the other categories.

**Table 1:**
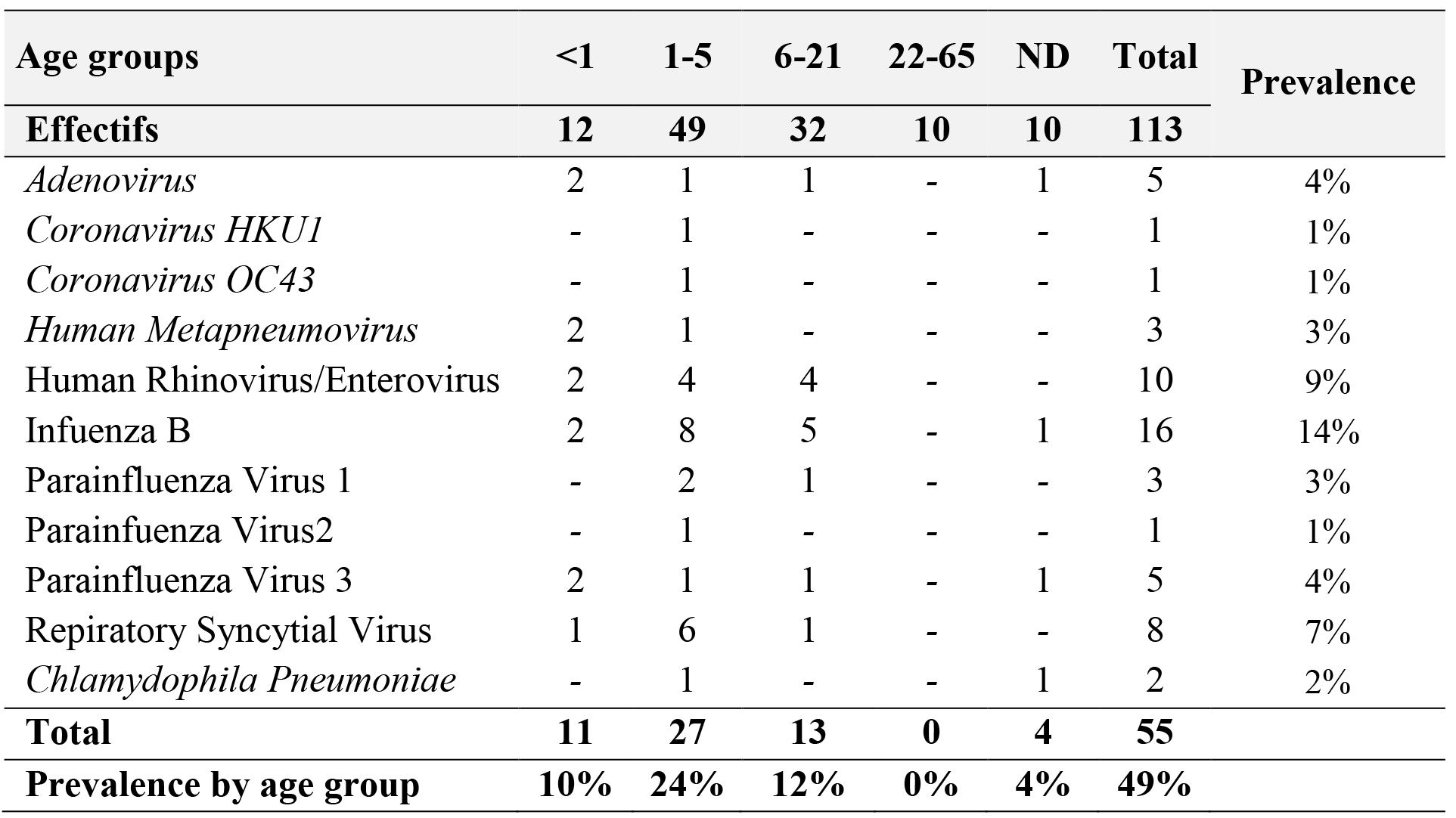
Representation of pathogens detected by age groups

Among the pathogens detected in winter, influenza B virus represented 36.4% (16/44) of cases and its prevalence during this period was about 20% (16/81). The difference between the prevalence rates during the dry and rainy seasons was not significant (χ^2^ = 1.93, *p*=0.16). The *Coronavirus* were only detected in the rainy season and the Influenza A virus was absent during the period of swabbing. The prevalence of the Influenza B virus was also significantly higher during the rainy season (Figure 3). The use of a respiratory multiplex panel allows the detection of 7 Co-infection (7/113, 6%) including adenovirus / human metapneumovirus, adenovirus / human rhinovirus, human rhinovirus / parainfluenzae virus, human rhinovirus / respiratory syncytial virus (n=2), influenza B / respiratory syncytial virus and parainfluenzae virus / *Chlamydia pneumoniae*.

**Figure 3:**
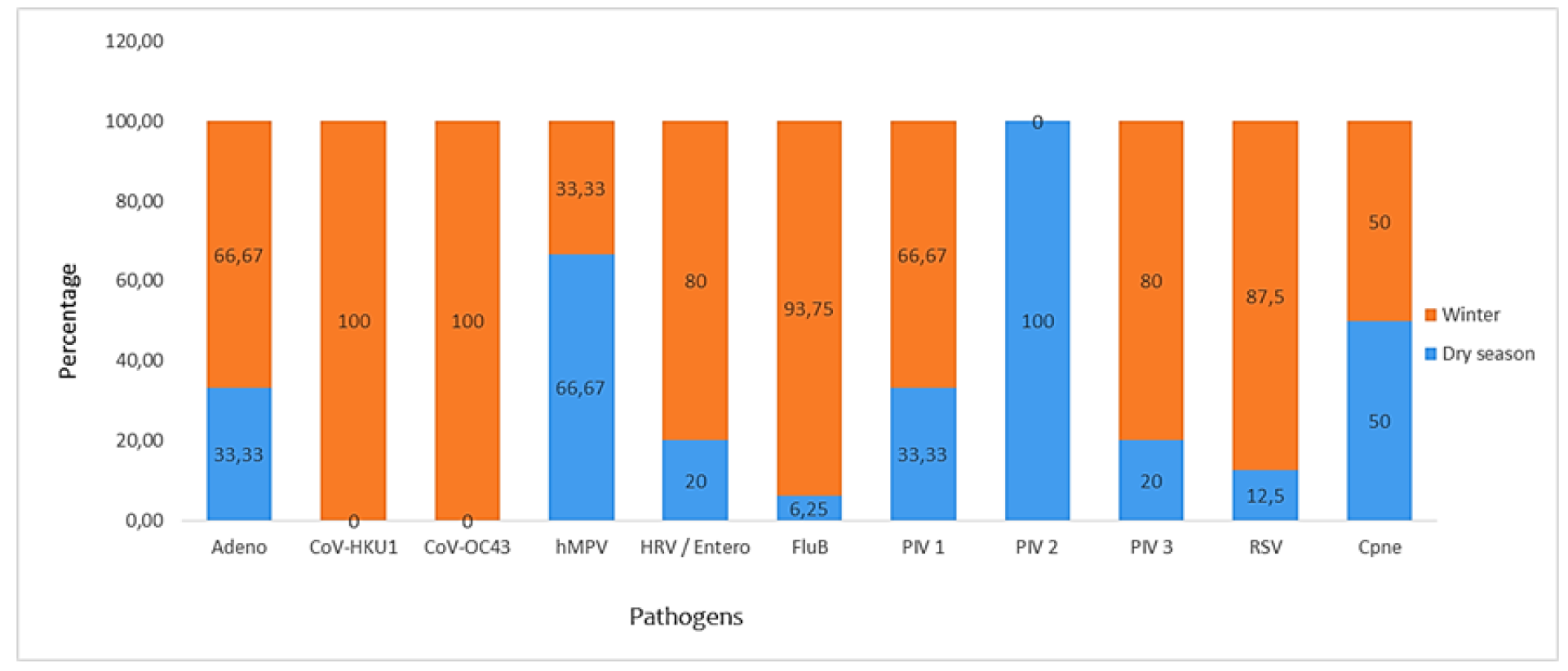
Seasonal prevalence of respiratory pathogens.

## Discussion

The diagnosis and subsequent management of respiratory tract infections is a real problem in developing countries, particularly for children under age 5 (2, 11). The situation is even more worrying for population of rural areas where the identification of pathogens responsible of infectious diseases is almost impossible. In this study, respiratory pathogens were identified in a high proportion (42%) of febrile patients with respiratory symptoms. The absence of negative controls represents a limit for this study, because we could not compare these results to those of subjects without signs and symptoms of respiratory tract infection in the same area. The analysis of samples from such subjects could provide information on the specificity of the detection by the FilmArray Respiratory Panel. But the results of week 37 of the year 2018, obtained by the 4S network (work focused on the surveillance of influenza viruses in Senegal) in the Fatick region, confirms the performance of FilmArray. To our knowledge, no data on the asymptomatic carriage of respiratory pathogens has been published in Senegal or Africa. Also, few studies focusing on the asymptomatic carriage of respiratory pathogens have been published in the world. However, some studies carried out in the USA report an asymptomatic carriage of 11% among health care personnel (12, 13). These persons are probably contaminated by the sick patients they care for. However, knowledge of the asymptomatic carriage rate is very important in the fight prospects. The disease surveillance and control strategy depends on reliable estimates of the proportion of asymptomatic carriers and the contribution of asymptomatic individuals to the transmission of respiratory pathogens. Pathogens were identified throughout the study, but the fact that the prevalence was higher during the rainy season suggests seasonality in the transmission of pathogens. However, in this study, the collection of specimens during a limited period constitutes a limit, because seasonality couldn’t be demonstrated. Moreover, the analysis by the FilmArray respiratory panel doesn’t show a significant difference between the prevalence of respiratory pathogens during the dry season and winter. Previous studies, performed in some Asian, European and African countries (Egypt and Senegal), demonstrated the relationship between climate and viral respiratory infections (14–19).

Results confirm that among the population, children under 5 years of age are more susceptible for respiratory infection (Table 1). For many infectious diseases, the 0-5 years of age group is more exposed than other groups. It should also be noted that the 22-65 age group was not diagnosed as being infected with tested pathogens. One of the explanations could be the low number of samples collected, which could result from the banalization of respiratory tract infections in this population, especially in rural areas. The same observations were made by Niang et *al*., 2012 who found that adults were less likely to consult for Influenza-like illness (20). Alternatively, this observation may be related to carriage of lower pathogen inoculums in adults than in children, as previously reported in other countries (21, 22). We believe that, in order to obtain the true prevalence of respiratory infections for this age group, studies based on active surveillance for respiratory tract infections are needed.

Concerning the pathogens identified, we noticed the absence of influenza A virus. This observation is confirmed by the report of the week 37-2017 of the 4S network, which performed a syndromic surveillance of infection diseases in Senegal. They notice the high prevalence of influenza B during this period and the absence of influenza A (23). The influenza B virus-like in Central African Republic (24), was the most commonly detected respiratory pathogen. It infected a high proportion of the population. Apart from the 22-65 years group, all the population was affected and especially the 1-5 years group. A recent study performed in Senegal highlighted the circulation of all lineages of influenza B virus in Senegal. They noticed that the circulation of Victoria lineage B virus intensified as the season progressed. The H1N1 pdm09 strain appeared midway and circulated extensively until the end of their study period. The circulation of H3N2 strain was negligible (19). The monitoring of influenza infection in 15 African countries performed from 2006 to 2010 revealed an overall increase of influenza-like illness (ILI) from 21 to 127 cases and severe acute respiratory infection (SARI) from 2 to 98 cases. Cases noticed in children between 0-4 years represented 48% of all ILI and SARI (25). In this study, the results show that children <5 years represented 48% of all ILI and 39% of all SARI. But the highest prevalence was noticed in children from 10-14 years. Unlike our results, they detected influenza virus in people >22 years. But the prevalence of influenza virus in ILI and SARI decreases as the age increases (25). Few data were published on the parainfluenza virus (PIV) burden in Africa. In Central African Republic data reported a prevalence of 3.3% for PIV-1 and 2.7% for PIV-3 (24). In Central African Republic as in Senegal, children under 5 years of age seem to be the most affected by the PIV. These results are similar to those observed in the United States of America where the rate of HPIV-1-associated hospitalization among children <5 years is estimated at 0.32–1.6 per 1000 children (26). The RSV is considered as the most common pathogen causing severe lower respiratory tract infections among infants and children (16). The prevalence of this pathogen was higher during the winter season but the difference observed in our results was not significant. We noticed seasonal differences in the apparition of this pathogen, as described by previous studies (16). As expected, RSV was essentially identified in the 0-5 years group, suggesting that this group is more susceptible to infection. This observation is very important, because the World Health Organization (WHO) is developing RSV surveillance (27) and closely following the RSV vaccine development (28). These results are similar to those observed in Central African Republic where a prevalence of 3.3% was noticed (24). But they are inferior to the results observed in Cameroon, Nigeria and Gabon where high prevalence rates of RSV were detected in febrile children with 7% in Cameroon (29), 17.7% in Nigeria (30) and 13.5% in Gabon (31). The study performed in Nigeria identified two subtypes of RSV, A and B, with a predominance of the subtype A. All the subtypes B identified in the study of Ibadan belonged to the BA genotype (30). These data would allow health authorities to target the specific population to be vaccinated. The FilmArray respiratory panel can’t make differentiate between adenovirus 1 and 2 and between the rhinovirus group and the enterovirus group.

The study has demonstrated an important burden of respiratory pathogens and the high prevalence of respiratory pathogens in the study area. The integration of the FilmArray respiratory panel into the field POC has been very beneficial to the population. Respiratory tract infections were diagnosed on site and results were obtained in a shorter amount of time. Health personnel were able to receive considerable support for the diagnosis of respiratory tract infections. Before the installation of BioFire, it was impossible to identify the respiratory pathogens circulating in this area, with the exception of influenza viruses diagnosed by RDTs. The people who consulted in the three health posts were able to benefit from adequate medical care with appropriate treatment. From a social point of view, improving the health of family members will save time for parent, which can then be used for farming or profit-making activities. The results of the FilmArray respiratory panel were similar to those from the Ministry of Health, published in week 37 in the Fatick region (23).

## Acknowledgements

The autors would like to thank bioMérieux that provided all the BioFire FilmArray^®^ material and the Respiratory Panel reagents that were used for the molecular diagnosis.

## Ethical approval

The project was approved by the National Ethics Committee N ° N° 00 098/MSP/DS/CNERS. Written informed consent was obtained from all individuals, including patients and parents or legal guardians of all children.

